# Multiscale conformational sampling of multidomain fusion proteins by a physics informed diffusion model

**DOI:** 10.64898/2026.03.11.711061

**Authors:** Zhaoqian Su, Bo Wang, Yinghao Wu

**Author notes:** Corresponding author: Yinghao Wu.

## Abstract

Multidomain fusion proteins, such as bispecific antibodies, rely on highly flexible linker regions for their therapeutic efficacy. Characterizing these vast conformational ensembles is crucial for rational drug design; however, while all-atom molecular dynamics (MD) is the traditional gold standard, its immense computational cost makes simulating large-scale domain motions prohibitive. Recently, deep generative diffusion models have emerged as a rapid alternative for sampling protein dynamics. Yet, being trained primarily on massive databases of structured, static domains, these generic models often lack the biophysical constraints required to thoroughly sample the large-scale dynamics of highly flexible multidomain architectures. To overcome this, we leverage microsecond MD trajectories of a multidomain protein construct with various linkers to train a multiscale diffusion framework utilizing an Equivariant Graph Neural Network (EGNN). To efficiently model the dynamics of the large molecular complexes, we employ a coarse-grained spatial graph that condenses rigid domains into center-of-mass anchors while preserving explicit backbone resolution for the flexible linker. By further integrating foundational rules in biophysics directly into both the training objective and the inference process, our model generates high-fidelity conformational ensembles that reproduce the thermodynamic distributions of long-timescale MD. This physics-informed approach provides a mathematically stable, highly scalable platform for the rapid multiscale characterization of flexible biologics, significantly accelerating the rational design of fusion protein therapeutics.

## Introduction

The biopharmaceutical industry is currently undergoing its fourth major revolution. This new wave of innovation is defined by multispecific drugs [1]. They offer profound clinical and biophysical advantages over their predecessors: while small molecules often struggle to disrupt large protein-protein interactions, conventional monoclonal antibodies are restricted to binding a single target. Among these next-generation modalities, fusion proteins have emerged as a uniquely powerful category [2]. Engineered by genetically fusing the coding sequences of two or more distinct functional proteins into a single open reading frame, these chimeric molecules are specifically designed to simultaneously engage multiple biological targets. In the context of modern immunotherapy, this dual-targeting capability allows fusion proteins to physically bridge distinct cellular environments [3]. Structurally, almost all of these engineered constructs share a common architectural motif: multiple rigid, structured functional domains connected by intrinsically disordered, flexible peptide linkers. Because of these unstructured linkers, these multi-domain constructs do not exist as static entities; rather, they undergo immense, continuous conformational dynamics [4]. The thermodynamic reach, spatial orientation, and overall physical flexibility governed by these linkers play an essential role in determining the target-binding kinetics and ultimate clinical efficacy of the multispecific therapeutic [5, 6]. Therefore, rigorously characterizing the dynamic conformational ensembles of these highly flexible constructs is of paramount importance for the rational design and optimization of next-generation biopharmaceuticals.

Computational methods serve as an effective approach to capture the dynamics of proteins. Among these tools, molecular dynamics (MD) remains the standard for capturing the temporal, physical interactions of biomolecules [7–14]. However, the immense computational cost of MD renders the systematic, long-timescale evaluation of large biomolecular libraries practically infeasible without dedicated supercomputing resources. To bypass this bottleneck, machine learning (ML) has emerged as a highly scalable alternative for characterizing protein dynamics. Fueled by the exponential expansion of the Protein Data Bank (PDB) [15], deep learning has already revolutionized static structural prediction, as demonstrated by AlphaFold [16, 17] and RoseTTAFold [18]. Building on this paradigm, researchers are increasingly leveraging MD trajectories to train ML models capable of predicting physically accurate conformational ensembles [19]. Furthermore, generative architectures, such as variational autoencoders and adversarial networks, have proven effective at compressing complex conformational landscapes into latent spaces, enabling the rapid sampling of novel structural states [20, 21]. Recently, generalized foundation models such as BioEmu have made significant impacts by learning latent representations directly from massive MD trajectory databases, enabling the rapid generation of diverse conformational states while bypassing traditional Newtonian integration steps [22]. However, these generalized foundation models have not been applied to large, highly flexible multi-domain fusion proteins that do not exist in PDB. The dynamics of these engineered constructs are governed by intrinsically disordered linkers and their resulting conformational landscapes are system-specific. Therefore, the development of a specialized generative model capable of efficiently and accurately sampling the unique conformational space of these flexible assemblies is critically needed.

To meet this need, we developed a multiscale computational framework that integrates long-timescale MD simulation with a physics-based diffusion model to capture the dynamics of a multispecific fusion protein. As a prototypical test system, we modeled a bispecific biologic comprising two natural immune receptor ligands, MHC and PD-L1, artificially connected by a flexible peptide linker. Analogous to natural naive T cell modulation, which requires both primary T cell receptor (TCR) engagement by MHC and a secondary coregulatory signal (**Figure 1a**) [23, 24], this MHC/PD-L1 conjugate is designed to selectively target and modulate only T cells expressing the corresponding TCR (**Figure 1b**). Because we hypothesize that the efficacy of this dual-receptor engagement is fundamentally dictated by the linker’s conformational flexibility, efficiently mapping these dynamics is critical. To achieve this without the prohibitive computational cost of traditional brute-force simulation, we trained a linker-conditioned Equivariant Graph Neural Network (EGNN) within a Denoising Diffusion Probabilistic Model (DDPM) architecture. By compressing the terminal folded domains into rigid bodies while preserving the flexible linker at higher resolution, we drastically accelerated conformational generation. We also applied a time-masked, physics-informed loss function to enforce proper bond lengths and angles. Our model generated structural ensembles that matched the macroscopic thermodynamics of MD simulation trajectories. Ultimately, this framework will unlock the rapid, high-throughput structural evaluation and thus accelerate the rational design of next-generation multispecific therapeutics.

**Figure 1:**
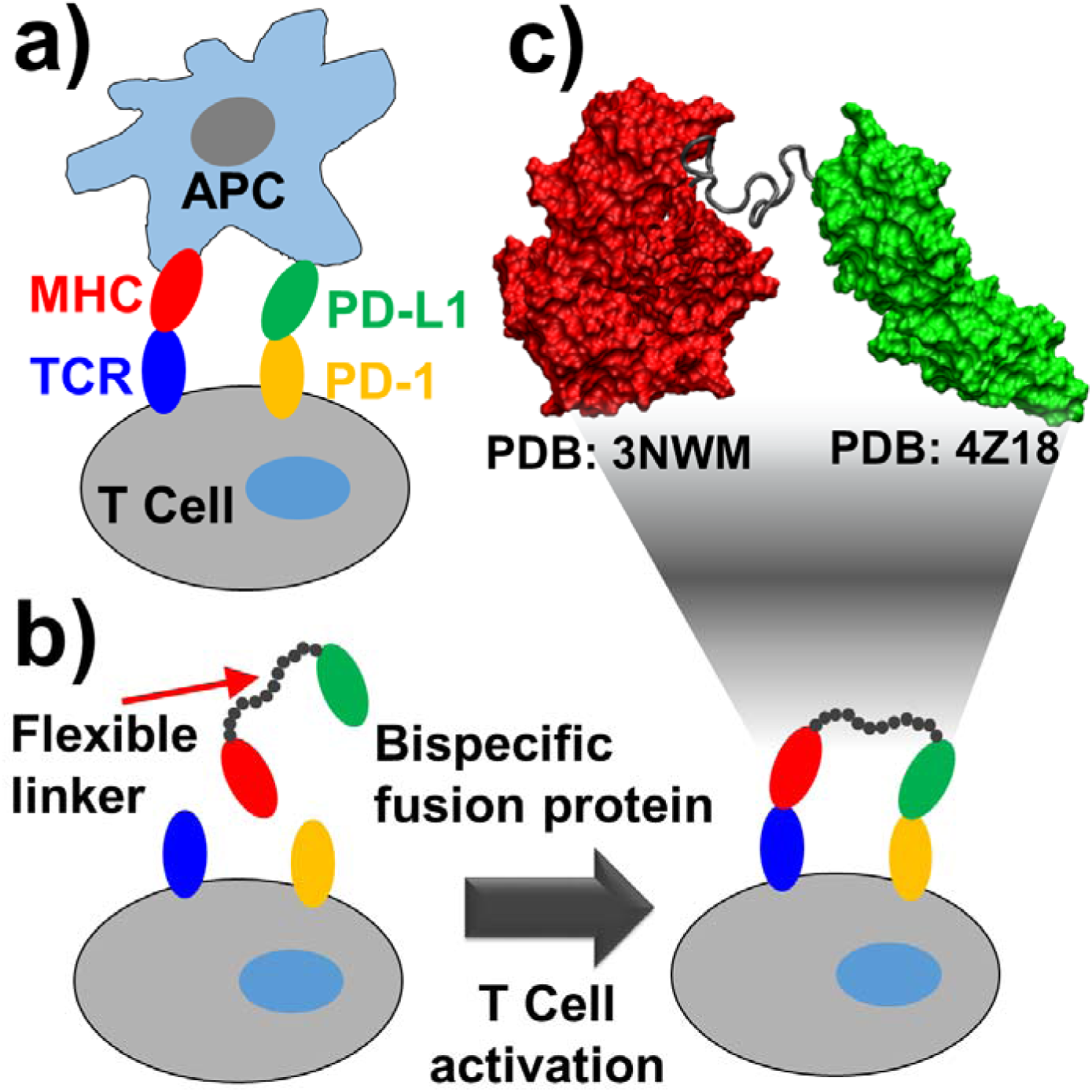
Biological context of bispecific fusion proteins. **(a)** Schematic of the MHC/TCR and PD-L1/PD-1 pathways modulating T-cell activation. **(b)** A bispecific therapeutic utilizes a flexible linker to simultaneously bridge distal receptors. **(c)** Structural model of the multidomain construct (MHC: red, PD-L1: green) connected by an intrinsically disordered flexible tether (gray).

## Methods

### System preparation and protocol of MD simulations

We computationally constructed a fusion protein by linking the MHC module (PDB: 3NWM) and the PD-L1 module (PDB: 4Z18) (**Figure 1c**). The 375-residue MHC module (H-2Kd heavy chain and β2m light chain) [25] was modeled without its groove peptide to ensure simulation stability. The PD-L1 module comprises two ∼100-residue immunoglobulin domains [26]. A peptide linker connects the MHC light chain C-terminus to the PD-L1 N-terminus. We designed two types of linkers: The first is called GS15 with a total length of 15 amino acids. The linker contains three repeats of small fragment with four glycine followed by a serine. The second is called GS30 with six copies of GGGGS fragments. These two linkers are assumed to be intrinsic disorder due to the flexible feature of glycine [27]. As a result, the initial structures of these two linkers were built by ModLoop [28]. Biologically, this bi-specific construct is designed to bypass the need for antigen-presenting cells (APCs): the MHC module targets specific T cell clones, while the PD-L1 module delivers a targeted co-regulatory inhibitory signal.

Equilibrium simulations (∼252,000 atoms per neutralized, water-solvated system) of this multi-domain protein complex were performed on the Anton 2 supercomputer [29]. Production runs utilized the NPT ensemble (1atm, 310K) with a Nosé-Hoover thermostat, a 2fs time step, and periodic boundary conditions in ∼125×120×160 Å orthorhombic boxes. We applied the CHARMM36m force field and the TIP4P-D water model [30], which improves the dynamic accuracy for flexible linkers [31]. Long-range electrostatics were computed using the Gaussian-split Ewald algorithm [32] on a 64×64×64 Å mesh, with an 11Å cutoff for short-range non-bonded interactions. Ultimately, a 2µs trajectory was collected and coordinates were saved every 1ns, yielding a total number of 2,000 dynamic snapshots for each molecular system.

### Data preprocessing and dimensionality reduction

To mitigate the immense computational complexity of all-atom ensembles, the molecular dynamics trajectories are mapped to a coarse-grained spatial graph. The structured domains of the complex, the MHC and PDL1, are mathematically collapsed into singular center-of-mass anchor nodes. In contrast, the highly flexible linker is preserved at explicit Cα backbone resolution to accurately capture its high-frequency conformational dynamics. This robust dimensionality reduction condenses the entire multi-domain fusion protein into a computationally efficient graph. Nodes are subsequently assigned categorical embeddings to differentiate rigid domain anchors from flexible linker beads, and spatial coordinates are geometrically normalized to stabilize the neural network’s variance schedule during diffusion training.

### Equivariant Graph Neural Network (EGNN) architecture for linker dynamics

To model the thermodynamic conformational space of the multidomain complex, the core generative engine employs a customized Equivariant Graph Neural Network (EGNN) architecture [33]. The system treats the protein complex as a fully connected spatial graph, where nodes represent either the coarse-grained rigid domain anchors or the highly flexible linker beads. The architecture initializes with distinct categorical embeddings for node types (rigid vs. flexible) and dynamically conditions these hidden states on both the specific linker system identity and the normalized continuous diffusion time step (*t*). The model passes these conditioned node features (*h*) and initial spatial coordinates (*x*) through four stacked EGNN layers.

Within each layer, the network strictly preserves *E(3)* rotational and translational equivariance. Edge messages are computed using a multi-layer perceptron (MLP) with SiLU activations, taking into account the hidden features of connecting nodes and their squared pairwise distances. Crucially, the coordinate update mechanism calculates a scalar "force" that physically translates the nodes. To ensure mathematical stability during the highly stochastic early stages of reverse diffusion, this predicted force is explicitly clamped within a small range and normalized by the total number of nodes in the specific graph. After updating the node hidden features via aggregated edge messages, the final layer outputs a residual spatial shift (Δ*x*), which represents the network’s prediction of the injected Gaussian noise (ε_θ_) that must be subtracted to denoise the structural ensemble.

### Physics-informed diffusion training

To train the EGNN to generate thermodynamically valid linker conformations, we utilized a Denoising Diffusion Probabilistic Model (DDPM) [34] framework augmented with a Physics-Informed Neural Network (PINN) objective [35] (**Figure 2a**). During the forward training phase, the ground-truth coarse-grained coordinates generated from MD simulations were iteratively corrupted by injecting Gaussian noise over a discrete Markov chain of 500 time steps. The noise variance is dictated by a linear schedule. The network was trained to predict the cumulative noise added at any given timestep, conditioned on the node types and specific linker system identity. Standard diffusion models rely solely on Mean Squared Error (MSE) for noise prediction. To embed rigorous biophysical constraints directly into the learning process, our training step calculates a dual-component loss:

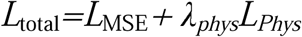

**Figure 2:**
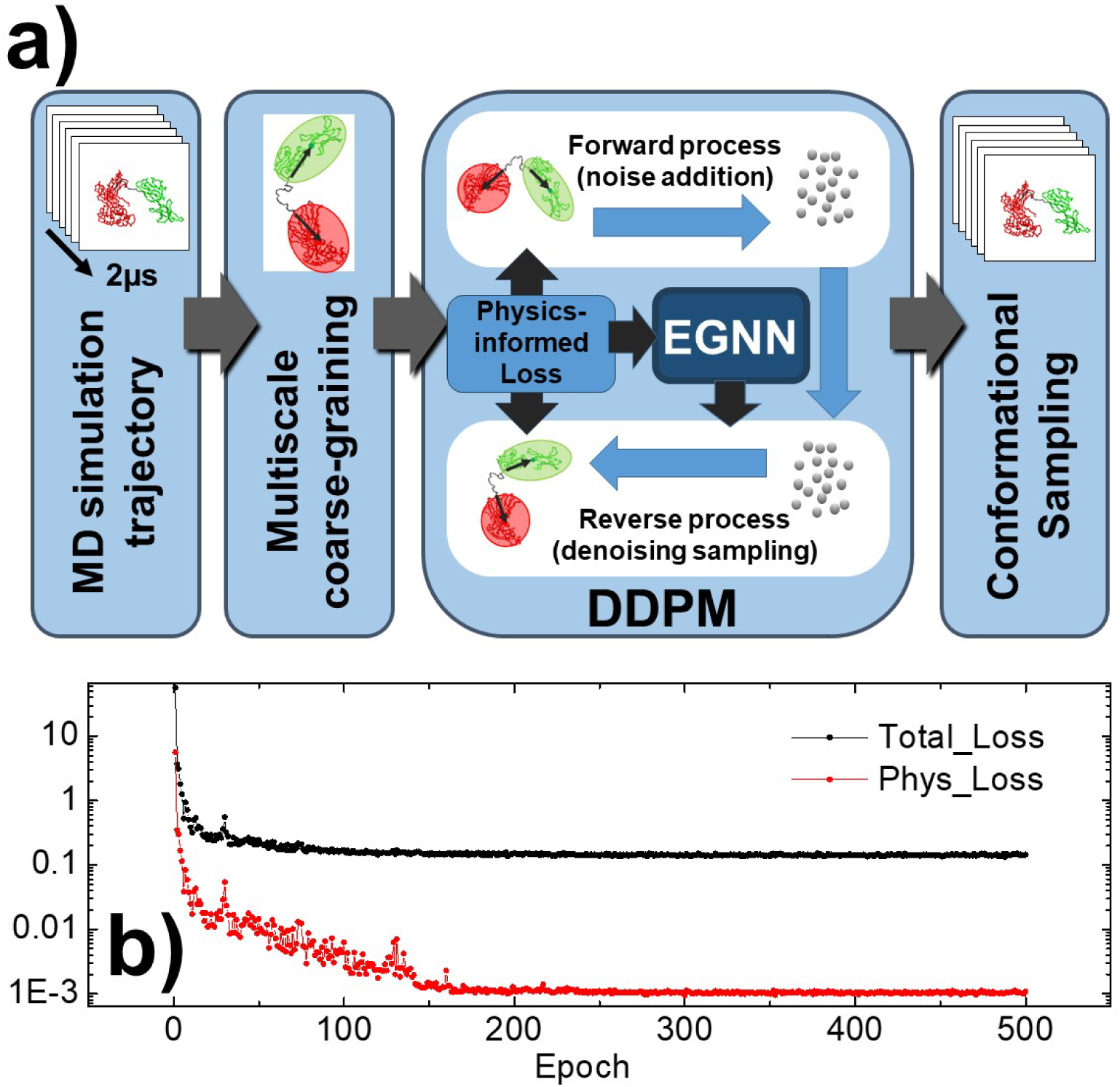
Physics-informed diffusion framework and training convergence. The schematic workflow of the framework is shown in **(a)**. MD trajectories are coarse-grained to train a diffusion model (DDPM/EGNN). A physics-informed loss function enforces local geometric constraints during the generative denoising process. **(b)** Training loss over 500 epochs. Total loss (black) converges rapidly, while the decoupled physics-informed loss (red) stably minimizes to enforce stereochemical accuracy.

The first term is the standard MSE between the true sampled Gaussian noise and the network’s predicted noise. The second term is to enforce continuous peptide backbone integrity. The model utilizes Tweedie’s formula to analytically estimate the fully denoised coordinates directly from the noisy state and the predicted noise.

The network is optimized using the Adam optimizer coupled with a *ReduceLROnPlateau* scheduler [36]. To prevent the rigid biophysical constraints from severely restricting the model’s ability to explore broad, global thermodynamic topologies, the physics penalty weight λ*_phys_*is dynamically annealed. Starting at an initial weight of 10.0, the penalty exponentially decays at a rate of 0.99 per epoch until it reaches a floor of 1.0. This allows the network to heavily prioritize localized structural integrity (bond lengths) in early epochs before seamlessly transitioning to optimizing the global conformational geometry of the entire fusion protein construct.

### Generative inference and structure reconstruction

To translate the trained diffusion model into biophysically meaningful structural ensembles, the inference pipeline couples the neural network’s thermodynamic predictions with strict geometric and kinematic algorithms. The inference process initializes the flexible linker as isotropic Gaussian noise and performs iterative reverse diffusion over 300 time steps to predict the macroscopic topology of the fusion protein. The raw generated coordinates are then translationally aligned to the static center-of-mass (CoM) of the reference MHC domain, anchoring the flexible linker to its correct biological origin. While the EGNN successfully captures the global free energy landscape, the stochastic nature of reverse diffusion can introduce microscopic, high-frequency spatial noise. To resolve this, the raw predictions are processed through a deterministic kinematic assembly layer. Starting from the MHC anchor, the algorithm sequentially reconstructs the linker bead-by-bead. Every step is strictly normalized to exact 3.8 Å, the length between two consecutive Cα atoms.

To prevent the generation of linkers with unphysical properties, the model evaluates the virtual bond angle among three consecutive Cα atoms. The cosine of this angle is mathematically restricted to a biologically validated window. If a predicted turn violates these boundaries, the algorithm orthogonally projects the vector onto the nearest valid limit before generating the next residue. Once the flexible linker is reconstructed, the PD-L1 will be anchored to the final linker bead. A rotation matrix, computed using the predicted center-of-mass vector of the PD-L1 domain, is then applied to orient the molecule according to the network’s thermodynamic prediction. Finally, the pipeline employs a comprehensive steric clash detector. If any non-bonded atomic pair falls below the coarse-grained van der Waals threshold of 3.5 Å, the conformation is flagged as unphysical and autonomously discarded.

### Data and code availability

All the source codes of these simulations are available for download at https://github.com/wujah/DiffusionAnton. The source codes of the diffusion model were written in Python3. The repository also contains the all-atom MD simulations used as input files of the model were derived from the Anton2. The program is free for academic users and works on a Linux platform.

## Results

We first evaluated the training stability and the effective integration of spatial constraints within our diffusion framework by monitoring the network’s loss components over 500 epochs (**Figure 2b**). The total loss (black curve) exhibits a rapid initial decrease, converging within the first 50 epochs. This indicates that the Equivariant Graph Neural Network (EGNN) quickly and efficiently learns the underlying reverse-diffusion denoising process from the multiscale coarse-grained trajectories. Crucially, the isolated physics-informed loss (red curve), which penalizes unphysical local geometries such as improper pseudo-bond lengths and steric clashes, demonstrates a continuous, stable decay, ultimately converging after approximately 200 epochs. This decoupled convergence profile confirms that the network successfully internalizes the strict polymer physics constraints without destabilizing the primary generative objective. The steady minimization of the physics loss ensures that the final model generates conformations that are both thermodynamically representative and stereochemically accurate.

Representative structures extracted from the model demonstrate the conformational heterogeneity (**Figure 3a**). The figure shows that we successfully sampled a wide array of relative domain orientations, ranging from highly extended, rod-like states to collapsed, interacting topologies. This structural diversity highlights the model’s ability to smoothly explore the vast conformational space dictated by the flexible, intrinsically disordered linker (gray beads) while preserving the distinct spatial boundaries of the functional rigid domains (green and red beads).

**Figure 3:**
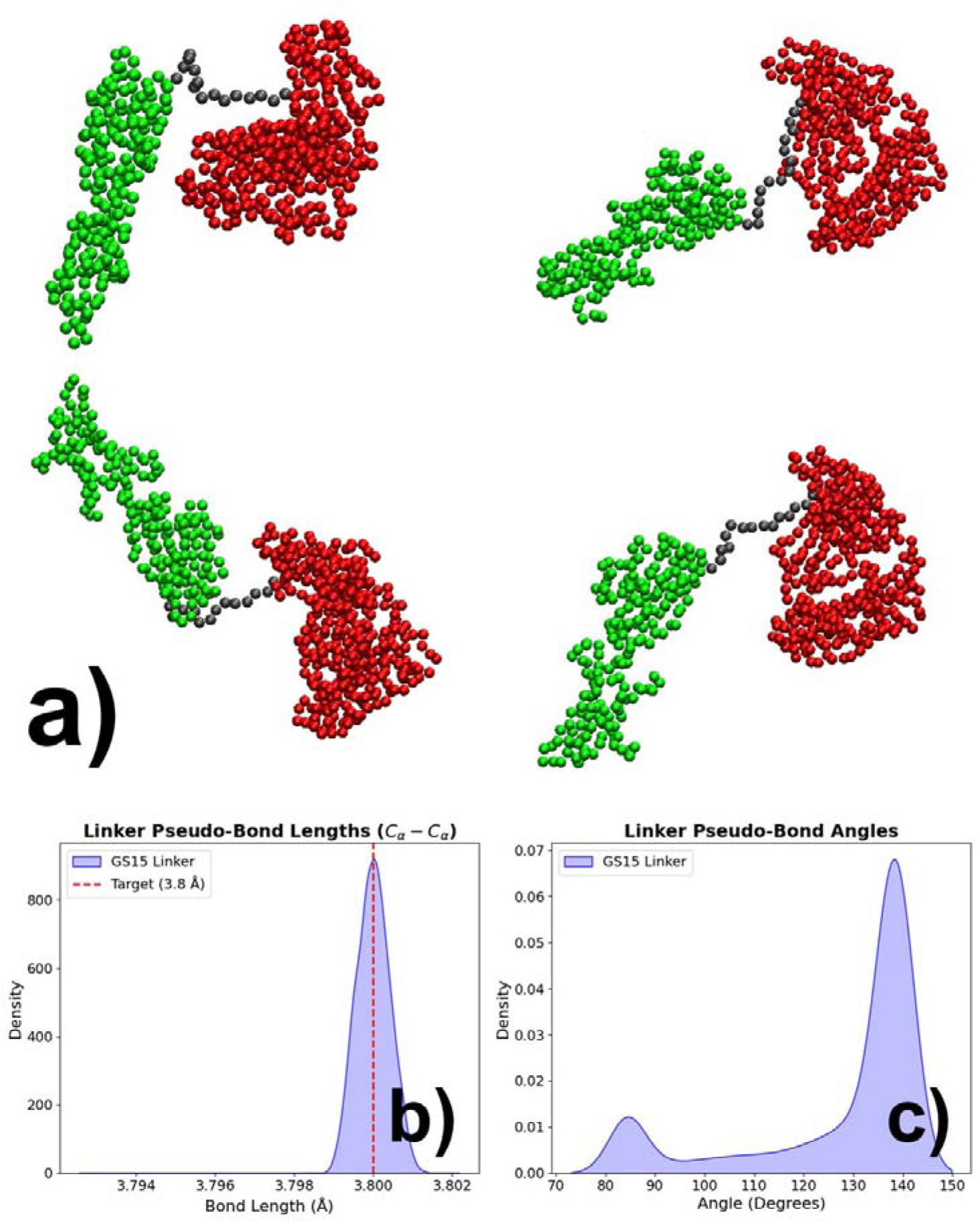
Structural diversity and geometric validation of the generated ensemble. **(a)** Representative coarse-grained structures illustrating the model’s ability to sample diverse compact and extended domain orientations. **(b)** The generated pseudo-bond length distribution strictly centers at the 3.8 Å target, confirming the prevention of unphysical chain stretching. **(c)** The pseudo-bond angle distribution exhibits realistic polymer flexibility while avoiding severe steric kinking.

Beyond macroscopic flexibility, a critical requirement for any generative structural surrogate is the strict preservation of proper local stereochemistry. We evaluated the internal coordinates of the generated linker to validate the efficacy of our physics-informed loss function. The distribution pseudo-bond lengths (**Figure 3b**) is centered precisely on the physically native target of 3.8 Angstrom. This confirms that the diffusion framework successfully enforced strict primary sequence connectivity, preventing the generation of unphysical chain breaks or artificial peptide stretching during rapid conformational sampling.

Furthermore, analysis of the pseudo-bond angles within the generated linker (**Figure 3c**) reveals a physically realistic profile. The distribution demonstrates natural polymer flexibility, defined by a major population centered near 135°–140° and a minor population near 85°. Crucially, the distribution sharply decays at lower angles, validating that the generative model inherently avoids severe steric clashes and unrealistic acute chain kinking.

Together, these geometric validations prove that the ensemble generated by the physics-informed diffusion model is not only thermodynamically accurate on a global scale but also highly robust at the local geometric level.

To further evaluate the accuracy of our model, we compared its structural ensembles against MD trajectories (**Figure 4**). **Figure 4a** shows that the model successfully captured macroscopic spatial constraints. The inter-domain distance distribution (MHC to PD-L1) of two ensembles well matched against each other, peaking symmetrically at ∼65 Å. Similarly, the linker’s localized compaction (radius of gyration, Rg) closely mirrored the native MD population near 10 Å, with the diffusion model sampling a slightly broader variance of extended states (**Figure 4b**).

**Figure 4:**
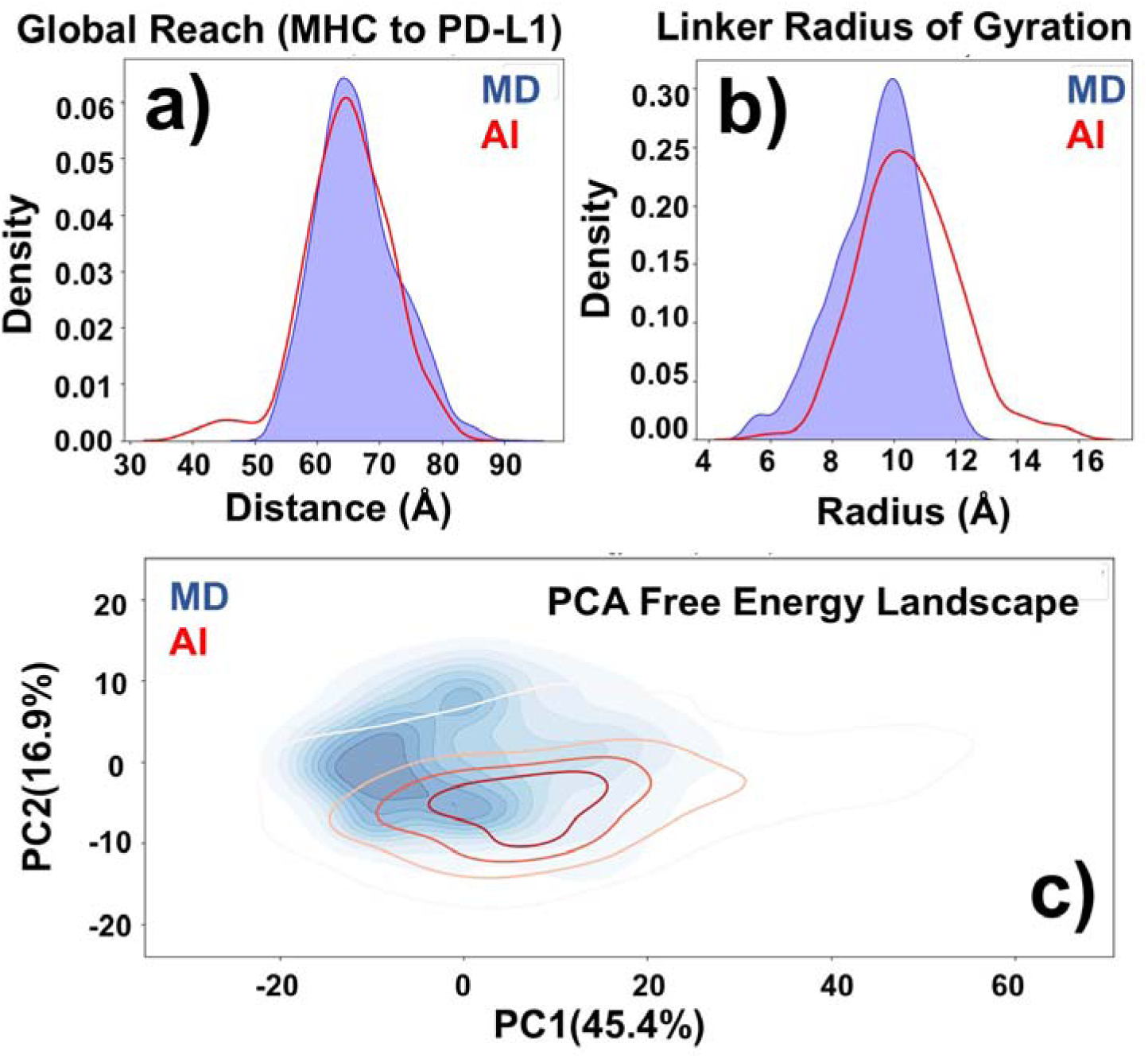
Validation of the model-generated ensemble against MD trajectory. Comparison of the generative (red) and MD (blue) models showing **(a)** global inter-domain distance and **(b)** linker radius of gyration, demonstrating high macroscopic agreement. **(c)** Projection onto a Principal Component Analysis (PCA) free energy landscape confirms the model accurately capture the native thermodynamic basins sampled by the MD simulation.

Furthermore, projecting both ensembles onto a shared Principal Component Analysis (PCA) free energy landscape confirmed that our model correctly navigates the multi-domain conformational space (**Figure 4c**). By reducing high-dimensional atomic fluctuations into a simple 2D topographical map, this landscape visualizes the protein’s dominant structural motions, where deep basins represent thermodynamically stable, low-energy states. The model-generated contours tightly overlapped these primary thermodynamic basins established by the MD simulation. This deep topological agreement proves our physics-informed diffusion framework not only accurately samples physically realistic, low-energy conformations, but also can effectively explores adjacent metastable states, without the prohibitive computational cost of microsecond-scale integration.

To evaluate the impact of linker sequence on the spatial dynamics of the bispecific biologic, we deployed our generative diffusion surrogate to sample the conformational ensembles of two MHC-PD-L1 fusion variants: a construct with a 15-residue linker (GS15) and a construct with a 30-residue linker (GS30). We quantified the structural heterogeneity of the generated ensembles by calculating the inter-domain distance (defined by the Center of Mass of the MHC and PD-L1 domains) and the global system Radius of Gyration (*R_g_*) across 250 fully reconstructed frames for each variant.

The inter-domain distance distributions reveal a stark contrast in the spatial reach afforded by the two linkers (**Figure 5a**). The GS15 construct exhibits a tightly constrained, sharply peaked distribution centered at approximately 65–70 Å, effectively acting as a rigid tether that prevents the functional domains from extending beyond 90 Å. In contrast, doubling the linker length to 30 residues (GS30) fundamentally alters the thermodynamic landscape. While the primary density peak shifts moderately outward to 80–85 Å, the GS30 ensemble develops a massive long-tail distribution extending past 160 Å. This pronounced tail indicates that the longer linker grants the bispecific construct the structural freedom to adopt highly extended, "open" conformations required to simultaneously bridge distal cellular receptors. Notably, the significant overlap between the two distributions in the 40–60 Å regime demonstrates that the GS30 variant retains the capacity to sample compact, collapsed states, highlighting a highly dynamic and continuous conformational space rather than a static extended rod. This localized spatial freedom translates directly into macroscopic structural heterogeneity, as captured by the overall system *R_g_* (**Figure 5b**). The GS15 ensemble remains conformationally restricted and largely globular, with its Rg sharply clustered around 40 Å. Conversely, the GS30 ensemble flattens and broadens considerably. The *R_g_* peak shifts to 45–48 Å, with the distribution populating a much wider basin that stretches up to 80 Å.

**Figure 5:**
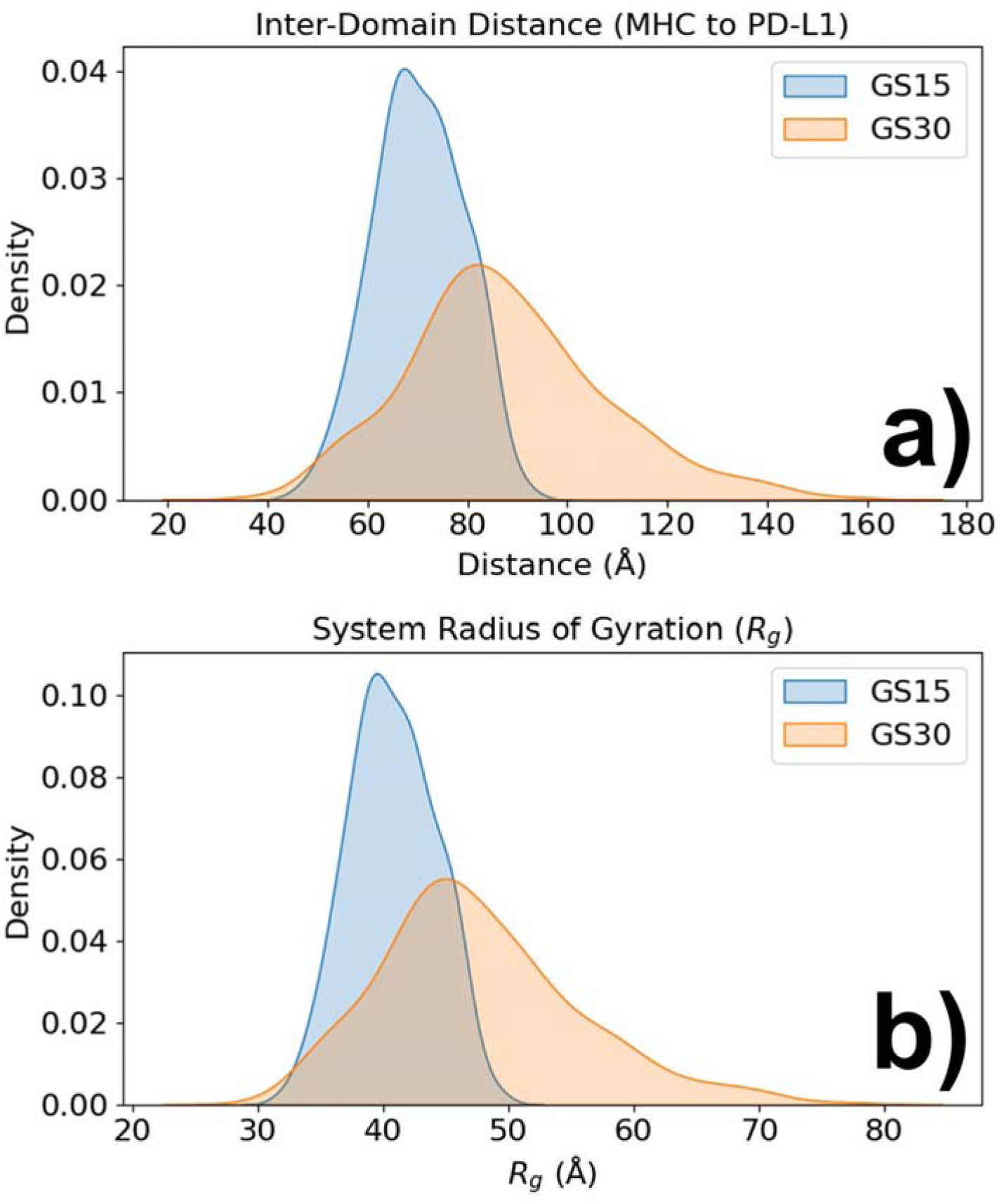
Impact of linker length on the thermodynamic ensemble. **(a)** Inter-domain distance reveals the longer GS30 linker provides a pronounced long-tail extension necessary for distal target engagement. **(b)** System *R_g_* distributions demonstrate that the GS30 linker introduces significant structural heterogeneity compared to the GS15 linker.

Together, these metrics successfully demonstrate that the intrinsically disordered linker actively governs the thermodynamic reach of the multi-domain biologic. Furthermore, the ability of our time-masked diffusion framework to rapidly and accurately capture this thermodynamic behavior without the prohibitive computational cost of microsecond-scale Newtonian integration, highlights its direct utility for the high-throughput, structure-based screening of next-generation multispecific therapeutics.

## Concluding Discussion

Multi-domain fusion proteins have emerged as highly promising therapeutic candidates across a broad spectrum of diseases, offering the unique ability to engage multiple distinct biological targets simultaneously. The pharmacological success of these multispecific biologics relies heavily on their macroscopic flexibility; the intrinsically disordered linkers must grant the rigid functional domains the precise spatial and thermodynamic reach required for simultaneous target engagement.

However, structure-based design for these highly flexible therapeutics is severely bottlenecked by current computational limitations. Traditional MD simulations are computationally prohibitive for adequately sampling the microsecond-to-millisecond conformational transitions required to evaluate such massive systems. Conversely, while state-of-the-art generative artificial intelligence models have revolutionized static protein structure prediction, they remain largely generic. Existing AI frameworks are not intrinsically tailored to predict the continuous thermodynamic ensembles of large, non-natural multi-domain proteins.

To bridge this gap, we introduced a physics-informed Denoising Diffusion Probabilistic Model (DDPM) utilizing a multiscale coarse-grained representation. By enforcing local geometric constraints during the generative denoising process, our framework successfully reproduces the native free energy landscapes of the MD ground truth at a fraction of the computational cost. Our results demonstrate that this generative model can rapidly and accurately predict how specific linker modifications alter the functional conformational ensemble of a multidomain biologic, providing a rigorously validated tool to accelerate next-generation drug design.

A fundamental challenge in applying generative AI to dynamic molecular ensembles is the notorious demand for massive training datasets. However, our framework demonstrates that a sparse dataset of 2000 MD frames is sufficient to train our model. This high data efficiency is directly driven by our multiscale coarse-grained representation. By abstracting the massive, structurally conserved functional domains (MHC and PD-L1) into rigid bodies and modeling the intrinsically disordered linker as a simplified Cα trace, we drastically reduce the system’s internal degrees of freedom. Consequently, the EGNN is relieved of the computationally immense burden of learning internal protein folding physics or high-frequency all-atom vibrations. Instead, the network focuses exclusively on the macroscopic translational and rotational dynamics of the domains and the polymer mechanics of the flexible tether.

Looking forward, the integration of physics-informed generative models into the biopharmaceutical pipeline offers a transformative approach to the structure-based design of flexible biologics. Because our multiscale DDPM framework effectively captures macroscopic thermodynamic ensembles at a fraction of the computational cost of traditional MD, it is uniquely positioned to enable the high-throughput in silico screening of therapeutic linker libraries. Rather than relying on empirical trial-and-error or prohibitive microsecond-scale simulations, researchers can rapidly generate and evaluate the precise spatial reach of hundreds of linker variants, assessing modifications in length, sequence composition, and rigidity in near real-time.

Furthermore, this multiscale methodology is highly extensible beyond the MHC-PD-L1 system presented here. The underlying framework can be readily adapted to model the conformational dynamics of other highly flexible, multi-domain architectures critical to modern pharmacology, such as bispecific T-cell engagers (BiTEs) [37], antibody-drug conjugates (ADCs) [38, 39], and targeted protein degraders like PROTACs [40, 41]. Ultimately, by providing a rapid, thermodynamically rigorous surrogate for conformational sampling, this approach alleviates one of the primary computational bottlenecks in rational biologic design, paving the way for the accelerated discovery and optimization of next-generation therapeutics.

## Acknowledgment

This work was supported by the National Institutes of Health under grant number R01GM120238, the United States–Israel Binational Science Foundation Project Number: 2023336, and the Einstein 2030 Seed Fund. The work was also partially supported by a start-up grant from Albert Einstein College of Medicine. Computational support was provided by Albert Einstein College of Medicine High Performance Computing Center. Anton 2 computer time was provided by the Pittsburgh Supercomputing Center (PSC) through grant R01GM116961 from the National Institutes of Health. The Anton 2 machine at PSC was generously made available by D.E. Shaw Research.

## Author Contributions

Z.S. and Y.W. designed the research; B.W. and Z.S. performed the research; Y.W. analyzed the data; Y.W. wrote the paper.

